# Gender-dependent effects of dietary oils on human PBMCs proliferation and redox status *in vitro*

**DOI:** 10.64898/2026.04.26.720862

**Authors:** Samia Bouamama

## Abstract

**Background:** Both dietary factors and biological sex are recognized as key modulators of immune responses. Nutritional components, particularly lipids, can influence immune cell metabolism, signaling pathways, and the balance between pro- and anti-inflammatory processes.

**Objective:** The present study aimed to examine whether commonly consumed dietary oils exert sex-specific effects on immune cell function and cellular oxidative balance.

**Methods:** Peripheral blood mononuclear cells (PBMCs) were isolated from 16 healthy adults (10 men and 6 women; mean age 48 years, BMI 23 kg/m^2^) using Histopaque density gradient centrifugation. Cells were cultured in RPMI-1640 medium and stimulated with concanavalin A (Con A) in the presence of olive, *Nigella sativa*, or walnut oils (23 μg/mL) for 48 h. Cell proliferation was assessed using the MTT assay. Intracellular malondialdehyde (MDA), protein carbonyls (PCAR), and reduced glutathione (GSH) were determined by spectrophotometric methods.

All statistical analyses were performed by Minitab 16 statistical software and Microsoft Excel 2007. Differences between groups were performed by Wilcoxon ranked test

**Results:** Baseline proliferation, MDA, and PCAR levels were comparable between sexes, whereas GSH levels were higher in male PBMCs. Oil supplementation significantly reduced proliferation in male cells compared to female cells (p = 0.008). In female PBMCs, olive oil significantly increased MDA levels, while all tested oils increased protein carbonyl levels. Walnut and olive oils selectively enhanced GSH levels in female cells.

**Conclusion:** Dietary oils modulate immune cell proliferation and oxidative balance in a sex-dependent manner. Female PBMCs appear more susceptible to lipid-induced oxidative stress, highlighting the importance of considering sex in nutritional immunology.

## 1. Introduction

The immune system plays a fundamental role in protecting the host against pathogens through complex cellular and molecular mechanisms. Its efficiency is influenced by intrinsic factors such as sex and extrinsic factors including diet [1].

Sex-related differences in immune responses are well established. Females generally exhibit stronger immune activation but are more prone to autoimmune diseases, whereas males are more susceptible to infections [2,3]. These differences are attributed to hormonal regulation and genetic factors, particularly those associated with the X chromosome.

Dietary lipids are key modulators of immune and oxidative processes. Fatty acids influence membrane composition, signaling pathways, and gene expression in immune cells [4]. Their biological effects depend on structural characteristics such as chain length and degree of unsaturation. Olive oil, rich in monounsaturated fatty acids, has recognized antioxidant and anti-inflammatory properties [5]. *Nigella sativa* oil contains polyunsaturated fatty acids and bioactive compounds with immunomodulatory potential [6]. Walnut oil is a major source of α-linolenic acid (omega-3), associated with cardioprotective and anti-inflammatory effects [7].

The present study aimed to evaluate the gender-dependent effects of these dietary oils from olive, *Nigella* or walnuts on PBMC proliferation and oxidative status *in vitro*.

## 2. Materials and Methods

### 2.1 Chemicals and Reagents

Chemicals used in this study were purchased from Sigma-Aldrich Company (Sigma, St Louis, MO, USA).

Incomplete Roswell Park Memorial Institute 1640 (RPMI 1640) medium was produced by Gibco Life Technologies Inc. (Paisley, U.K). it was aseptically supplemented with 25 mM HEPES buffer, 10% heat-inactivated fetal calf serum, L-glutamine (2 mM), 2-mercaptoethanol (5 × 10^-5^ M), penicillin (100 UI/ml) and streptomycin (100 μg/ml).

Each form of studied oils (olive, black seeds, walnut) was dissolved in ethanol 70%. The final concentration of oil solution in culture medium was 23 μg/ml, with a nontoxic concentration of ethanol (0.015 %).

### 2.2 Study Population

PBMCs were isolated from venous EDTA-blood freshly obtained from 16 healthy volunteers (10 men and 6 women). All participants provided informed consent, and the study was approved by the ethics committee of Tlemcen university hospital in accordance with the Declaration of Helsinki.

### 2.3 PBMC Isolation and Culture

PBMCs were isolated using Histopaque-1077 density gradient centrifugation. Cells were cultured in RPMI-1640 medium supplemented with fetal calf serum, antibiotics, glutamine, and 2-mercaptoethanol. Cell viability was confirmed using Trypan blue exclusion.

Cells (4 × 10□/well) were stimulated with Con A (5 μg/mL) and treated with olive, *Nigella sativa*, or walnut oils (23 μg/mL). Ethanol (0.015%) was used as vehicle control. Cultures were incubated for 48 h at 37°C in 5% CO□.

### 2.4 Cell Proliferation Assay

Cell proliferation was assessed using the MTT assay as previously described [8]. Briefly, after 48 h of incubation cell viability was assessed by adding 10 μl/well of MTT (5 mg/ml), the plates were read on micro-plates reader at a 570 nm wavelength. PBMCs Proliferation is expressed as a percentage of absorbance of treated cells to untreated control cells.

Or **Cell viability % =** {OD treated cells/ OD untreated cells controls × 100}

### 2.5 Intracellular oxidative stress markers

Oxidative stress biomarkers in PBMCs (Malondialdehydes: MDA, Protein carbonyls: PCAR, and reduced glutathione: GSH) were assessed by measuring their concentrations in cell homogenate.

MDA levels were measured using the TBARS assay [9]

Protein carbonyls were determined using the DNPH method [10]

GSH levels were quantified using Ellman’s reagent [11]

### 2.6 Statistical Analysis

All statistical analyses were performed by Minitab 16 statistical software and Microsoft Excel 2007. Data are expressed as mean ± SEM. Differences between groups was performed by Wilcoxon ranked test . All *in vitro* cultures were repeated at least three times. Statistical significance was set at *p*□< □0.01.

## 3. Results

Our results demonstrate that cell proliferation rate, intracellular MDA and PCAR levels were comparable between males and females while intracellular GSH levels were higher in male’s PBMCs non treated cells as compared to females.

It is worth to note that all tested oils diminished considerably cellular proliferation in males compared to females (*P*=0,008). Fig 1

**Fig 1.**
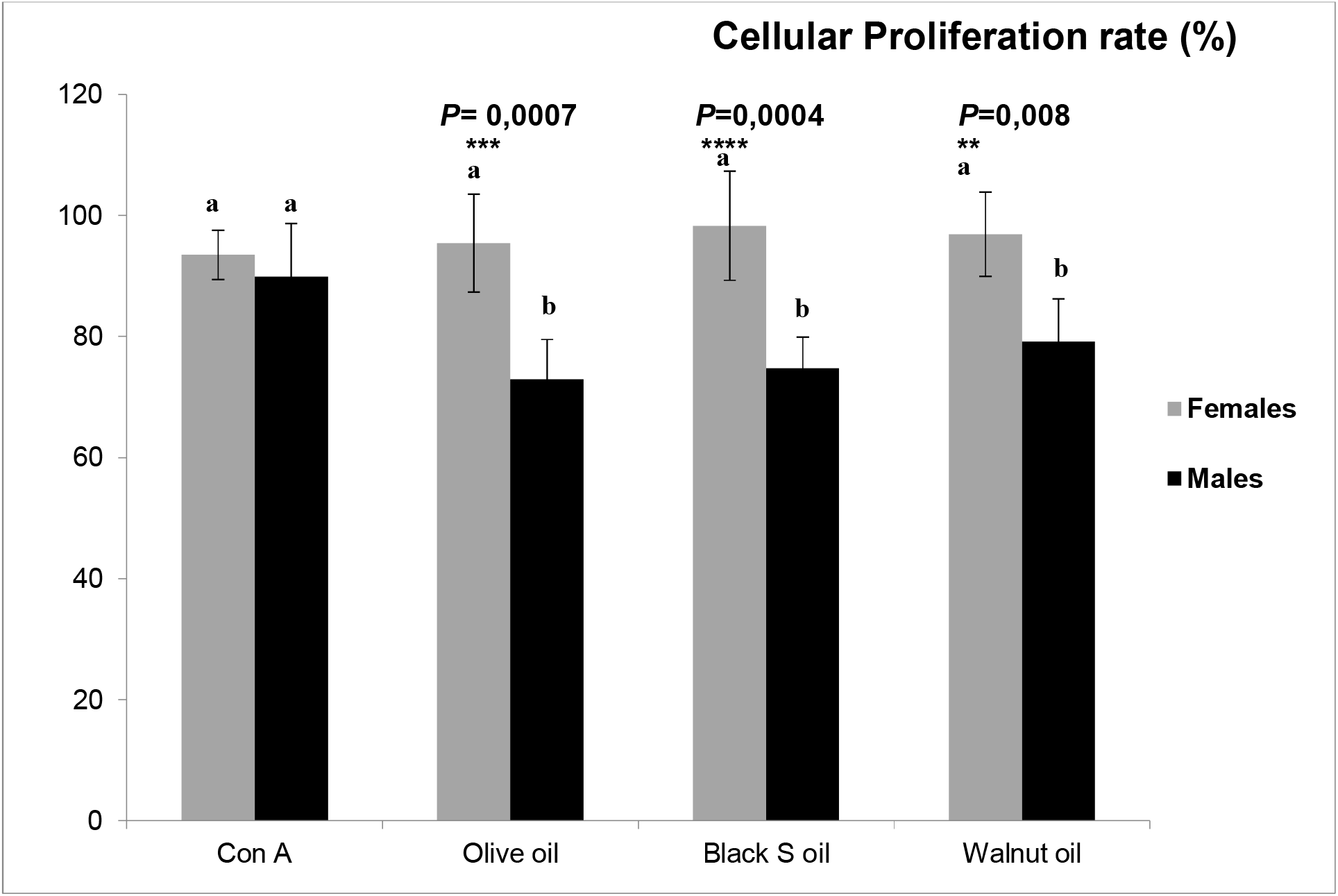
PBMCs cellular proliferation rate in response to dietary oils and Con a treatments. Each value represents the mean ± SEM. Significant differences between groups are indicated by small letters (a,b,c,d), with p less than 0.01. In the presence of olive oil, females cellular MDA levels increased significantly (*P*=0,008). Fig 2

**Fig 2.**
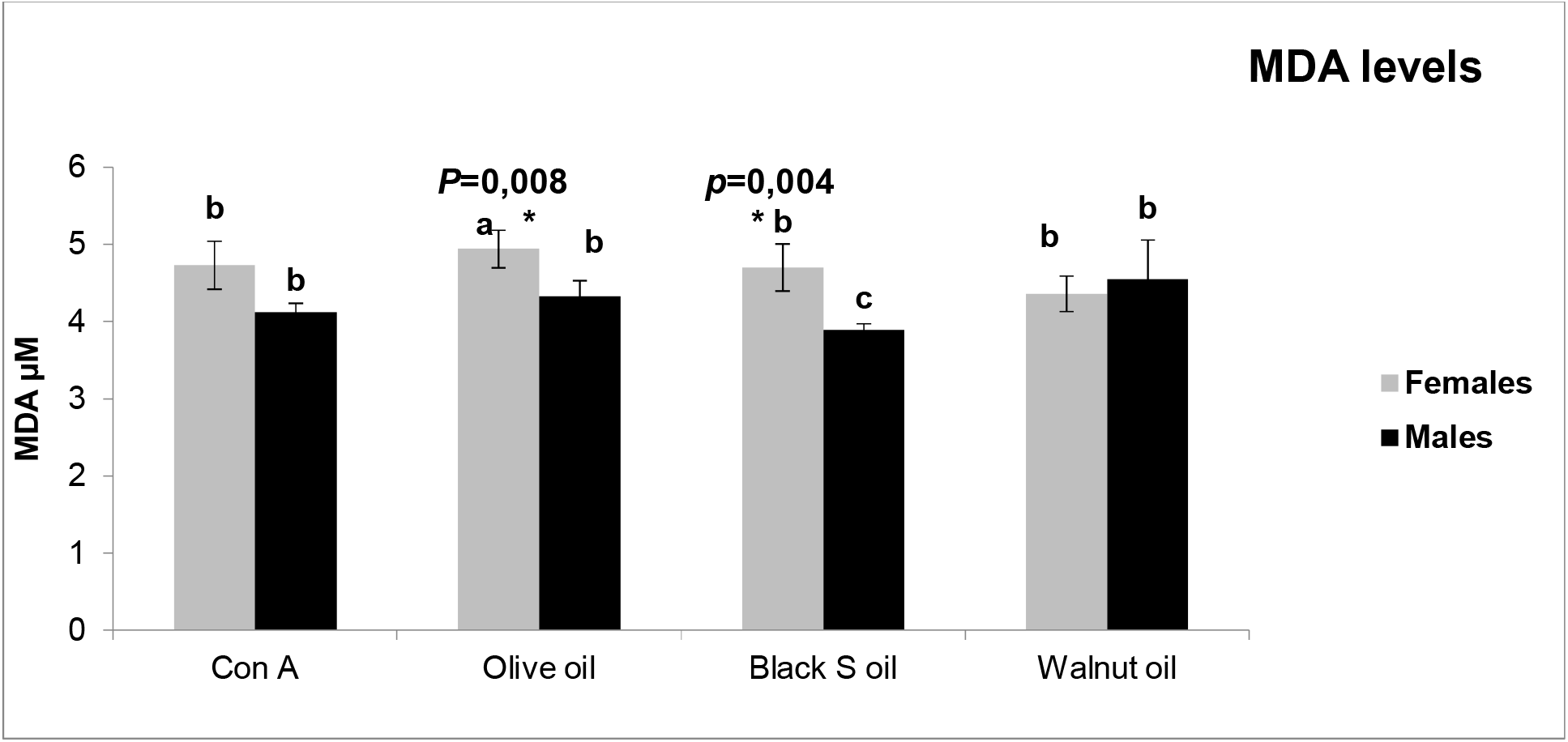
PBMCs cellular MDA levels in response to dietary oils and Con a treatments. Each value represents the mean ± SEM. Significant differences between groups are indicated by small letters (a,b,c,d), with p less than 0.01 PCAR levels increased significantly in female cells treated with all forms of studied oils, by contrast those of males are not affected. Fig 3

**Fig 3.**
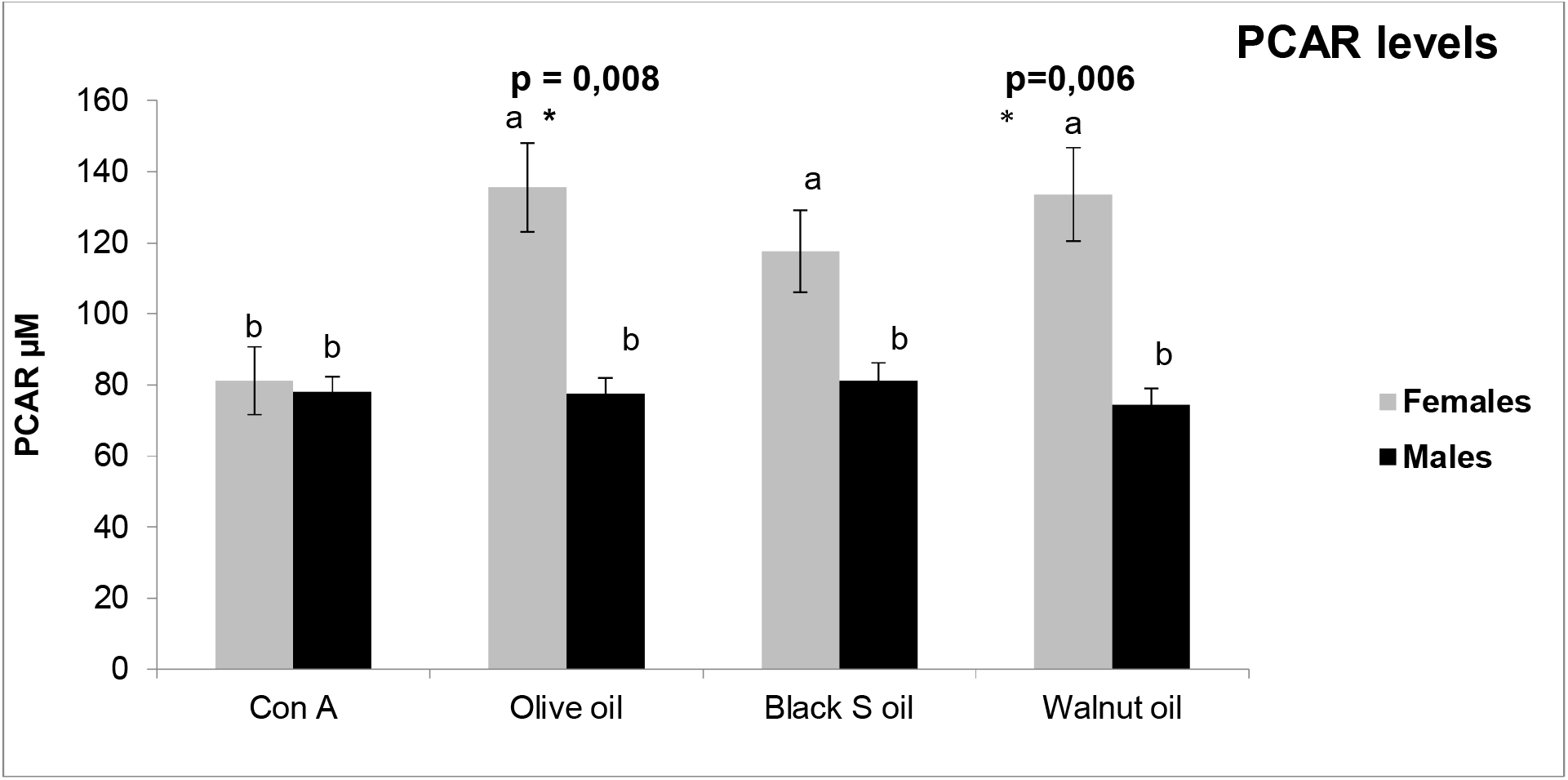
PBMCs cellular PCAR levels in response to dietary oils and Con a treatments. Each value represents the mean ± SEM. Significant differences between groups are indicated by small letters (a,b,c,d), with p less than 0.01. Concerning GSH levels, our results show that basic GSH levels in women are diminished compared to men (*p*= 0, 0008). Walnut and olive oil and not black seeds oil increased GSH levels only in females PBMCs. Fig 4

**Fig 4.**
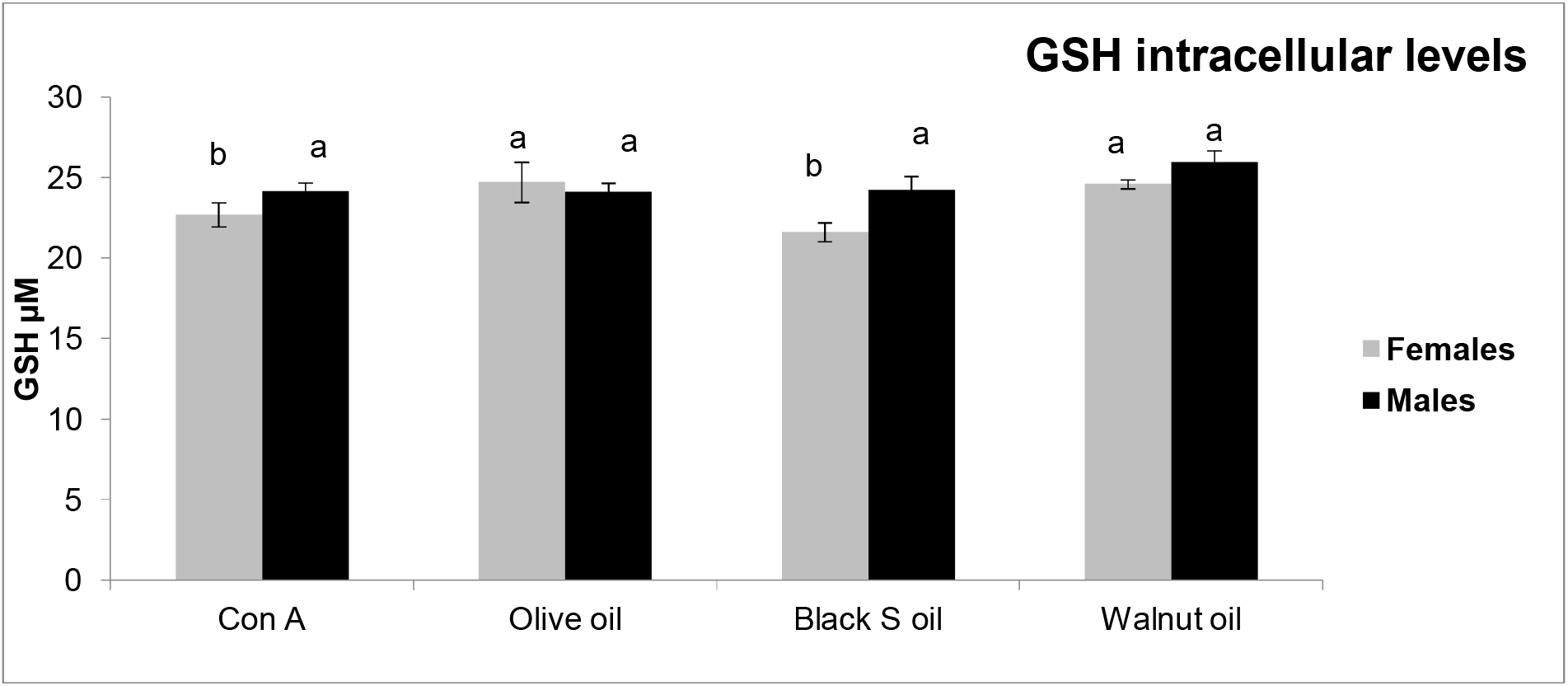
PBMCs cellular GSH levels in response to dietary oils and Con a treatment. Each value represents the mean ± SEM. Significant differences between groups are indicated by small letters (a,b,c,d), with p less than 0.01

## 4. Discussion

This study demonstrates that dietary oils exert sex-specific effects on immune cell function and redox balance. While basal immune parameters were comparable, responses to lipid exposure differed significantly between males and females.

The increased oxidative stress observed in female PBMCs is consistent with the higher susceptibility of polyunsaturated fatty acids to lipid peroxidation [12]. Fatty acids can also enhance ROS production via NADPH oxidase activation [13], contributing to oxidative damage reflected by increased MDA and protein carbonyl levels [14]. Sex-dependent differences may be explained by variations in immunometabolism. Female immune cells exhibit higher metabolic activity and immune responsiveness [15,2], which may increase sensitivity to oxidative stress. Additionally, mitochondrial differences may contribute to enhanced ROS production in female cells [16,17].

Hormonal factors also play a critical role. Estrogens regulate lipid metabolism and antioxidant defenses but may enhance oxidative stress under certain conditions [18,19].

The increase in protein carbonyls in female cells suggests altered immune function, as protein oxidation affects signaling and proliferation [14,20]. The observed increase in GSH levels in females likely represents a compensatory antioxidant response [19,17].

Fatty acids also modulate immune signaling pathways. Omega-3 and omega-6 fatty acids influence inflammation and immune responses in a sex-dependent manner [21,22].

Overall, these findings confirm that lipid-induced modulation of immune function is strongly influenced by sex and involves complex interactions between metabolism, hormonal regulation, and redox homeostasis.

## 5. Conclusion

Dietary oils modulate immune cell proliferation and oxidative stress in a sex-dependent manner. Female PBMCs demonstrate increased vulnerability to oxidative challenges induced by fatty acids, highlighting the need for sex-specific considerations in nutritional research and dietary guidance. Integrating sex as a critical variable can enhance our understanding of nutritional immunology and support the development of more tailored and effective dietary strategies.

## Acknowledgements

The author thanks all volunteers who participated in this study.

## References

1- Miles EA, Calder PC, Carr AC, et al. Diet and immune function. Nutrients. 2021;13 (1):290. doi:10.3390/nu13010290

2- Klein SL, Flanagan KL. Sex differences in immune responses. Nat Rev Immunol. 2016;16(10):626–638. doi:10.1038/nri.2016.90

3- Fransen F, van Beek AA, Borghuis T, Aidy SE, Hugenholtz F, van der Gaast-de Jongh C, et al. The impact of gut microbiota on gender-specific differences in immunity. Front Immunol. 2017;8:754. doi:10.3389/fimmu.2017.00754

4- Calder PC. Nutrition, immunity and COVID-19. BMJ Nutr Prev Health. 2020;3(1):74–92. doi:10.1136/bmjnph-2020-000085

5- Djelti F, et al. Olive oil components and their beneficial effects on human health. J Food Biochem. 2014;38(5):461–472.

6- Ahmad A, Husain A, Mujeeb M, Khan SA, Najmi AK, Siddique NA, et al. A review on therapeutic potential of Nigella sativa: A miracle herb. Asian Pac J Trop Biomed. 2013;3(5):337–352. doi:10.1016/S2221-1691(13)60075-1

7- Ros E. Health benefits of nut consumption. Nutrients. 2010;2(7):652–682. doi:10.3390/nu2070652

8- Mosmann T. Rapid colorimetric assay for cellular growth and survival: Application to proliferation and cytotoxicity assays. J Immunol Methods. 1983;65(1–2):55–63. doi:10.1016/0022-1759(83)90303-4

9- Nourooz-Zadeh J, Tajaddini-Sarmadi J, Wolff SP. Measurement of plasma lipid peroxidation products by thiobarbituric acid assay. Clin Chem. 1996;42(6):861–866.

10- Levine RL, Garland D, Oliver CN, Amici A, Climent I, Lenz AG, et al. Determination of carbonyl content in oxidatively modified proteins. Methods Enzymol. 1990;186:464–478. doi:10.1016/0076-6879(90)86141-H

11- Ellman GL. Tissue sulfhydryl groups. Arch Biochem Biophys. 1959;82(1):70–77. doi:10.1016/0003-9861(59)90090-6

12- Höhn A, Jung T, Grimm S, Grune T. Lipid peroxidation-derived aldehydes and their role in aging and age-related diseases. Mol Aspects Med. 2013;34(2–3):539–548. doi:10.1016/j.mam.2012.08.008

13- Cury-Boaventura MF, Curi R. Regulation of reactive oxygen species (ROS) production by fatty acids. Clin Sci (Lond). 2005;108(3):245–252. doi:10.1042/CS20040273

14- Dalle-Donne I, Rossi R, Giustarini D, Milzani A, Colombo R. Protein carbonyl groups as biomarkers of oxidative stress. Trends Mol Med. 2006;12(12):589–598. doi:10.1016/j.molmed.2006.10.001

15- Márquez EJ, Chung CH, Marches R, Rossi RJ, Nehar-Belaid D, Eroglu A, et al. Sexual-dimorphism in human immune system aging. Nat Commun. 2020;11:751. doi:10.1038/s41467-020-14396-9

16- Ventura-Clapier R, Dworatzek E, Seeland U, Kararigas G, Arnal JF. Sex in basic research: Concepts in the cardiovascular field. Cardiovasc Res. 2017;113(7):711–724. doi:10.1093/cvr/cvx066

17- Tower J. Sex differences in aging and oxidative stress. Antioxidants (Basel). 2023;12(6):1255. doi:10.3390/antiox12061255

18- Mauvais-Jarvis F. Aging, male sex, obesity, and metabolic inflammation create the perfect storm for COVID-19. Diabetes. 2020;69(9):1857–1863. doi:10.2337/dbi19-0023

19- Rubio-Ruiz ME, Guarner-Lans V, Pérez-Torres I, Soto ME. Mechanisms underlying metabolic syndrome-related oxidative stress. Oxid Med Cell Longev. 2019;2019:3640815. doi:10.1155/2019/3640815

20- Conti P, Caraffa A, Gallenga CE, Ross R, Kritas SK, Frydas I, et al. The role of inflammation and oxidative stress in the pathogenesis of COVID-19. Int J Mol Sci. 2020;21(21):8118. doi:10.3390/ijms21218118

21- Calder PC. Omega-3 fatty acids and inflammatory processes: From molecules to man. Biochem Soc Trans. 2023;51(1):1–14. doi:10.1042/BST20220152

22- Dai X, Ahn KS, Kim C, Siveen KS. Polyunsaturated fatty acids and their immunomodulatory roles in health and disease. J Funct Foods. 2021;85:104629. doi:10.1016/j.jff.2021.104629

